# Natural statistics of host odours predict species-specific olfactory behaviours in Drosophilids

**DOI:** 10.64898/2026.03.27.714575

**Authors:** Hui Gong, Zofia Ziolkowska, Mohammed A. Khallaf, Sinziana Pop, Oscar Ayrton, Xavier Cano-Ferrer, James MacRae, Markus Knaden, Roman Arguello, Lucia L. Prieto-Godino

## Abstract

Animals rely on olfaction to locate food, mates, and suitable habitats, yet natural odour environments contain thousands of volatile molecules, creating a high-dimensional sensory problem for both nervous systems and the researchers who study them ^1–5^. For example, a banana emits around 100 individual volatiles^4,6^. It remains unclear which components of complex odour blends animals have evolved to use as behavioural cues. Here, combining fieldwork, chemical and behavioural analyses, we show across multiple *Drosophila* species that behaviourally relevant cues can be predicted directly from the statistical structure of natural odour environments. Animals preferentially respond to components that are most distinctive within their natural host odour blends, and therefore most ecologically informative. These cues can be either major or minor blend components. Our results indicate that host-guided olfactory behaviours have evolved to exploit the statistical structure of natural odour environments by selectively targeting the most informative features of odour blends.

## Results and Discussion

Inspired by efficient coding theory, we asked whether behaviourally relevant cues could be predicted directly from the statistical structure of natural odour environments. If olfactory systems efficiently encode ecological information, behaviourally relevant cues should correspond to those odorants that carry the most information about resource identity. Such information is concentrated in odour compounds that consistently distinguish hosts’ odour mixtures. To test this, we analysed an existing GC-MS dataset of volatiles emitted by 34 fruit hosts of *Drosophila melanogaster*, comprising 2,491 detected compounds^4^ (Fig. 1a). Absolute GC-MS peak areas do not directly reflect sensory input, as early olfactory processing steps, such as gain control and lateral inhibition, emphasise relative contrast within odour blends. Therefore, to identify ecologically informative components, we transformed the data into a saliency normalised space (Methods). This transformation enables us to analyse fruits’ chemical profiles based on which odorants stand out relative to each fruit’s background bouquet, consistent with recent work showing that natural odour spaces are structured by the statistics of odorant co-occurrence in natural mixtures rather than by independent variation of individual compounds^2^. We then applied principal component analysis (PCA) to this normalized dataset (Methods). We used the loadings of the first three principal components to rank odorants that best distinguished fruits. Remarkably, variance along the first three PCs was primarily driven by only eight of the 2,491 individual compounds (Fig. 1b and Table S1). Furthermore, despite the analysis relying solely on the statistical structure of natural odour variation and being entirely agnostic to behavioural data, seven of these eight identified odorants had previously been shown to be behaviourally and ecologically relevant to *D. melanogaster*^4–11^ (Fig. 1b, Table S2). For example, D-limonene guides female oviposition choice^7^ and isoamyl acetate is a potent feeding stimulant for adult *D. melanogaster*^11^. Thus, natural statistics of host odours alone recovered known behaviourally active cues. This supports the hypothesis that informative compounds guide behaviour, leading us to predict that odours ranking higher in our analysis should be preferentially associated with attraction. To test this prediction systematically, we annotated published behavioural responses for 45 odorants (Table S2), and compared these to a “PC score”, a variance explained weighted average of the first three PC loadings (Fig. 1c, Methods). We found that behaviourally attractive compounds displayed significantly higher scores than neutral or ambiguous compounds (Fig. 1d). Therefore, compounds that carry more information about fruit identity are preferentially associated with attraction. Importantly, repeating the analysis after standardising variables across samples - a standard preprocessing step in PCA - or after centered log-ratio (CLR) normalization, a transformation recommended for analysing proportional chemical ecology datasets^12^, did not equally prioritise behaviourally relevant odorants (Fig. S1, Methods), indicating that accurate behavioural prediction depends on preserving within-blend contrast structure. Together, these analyses indicate that *D. melanogaster* olfactory host seeking behaviours are biased towards the most informative host fruit odours.

**Figure 1.**
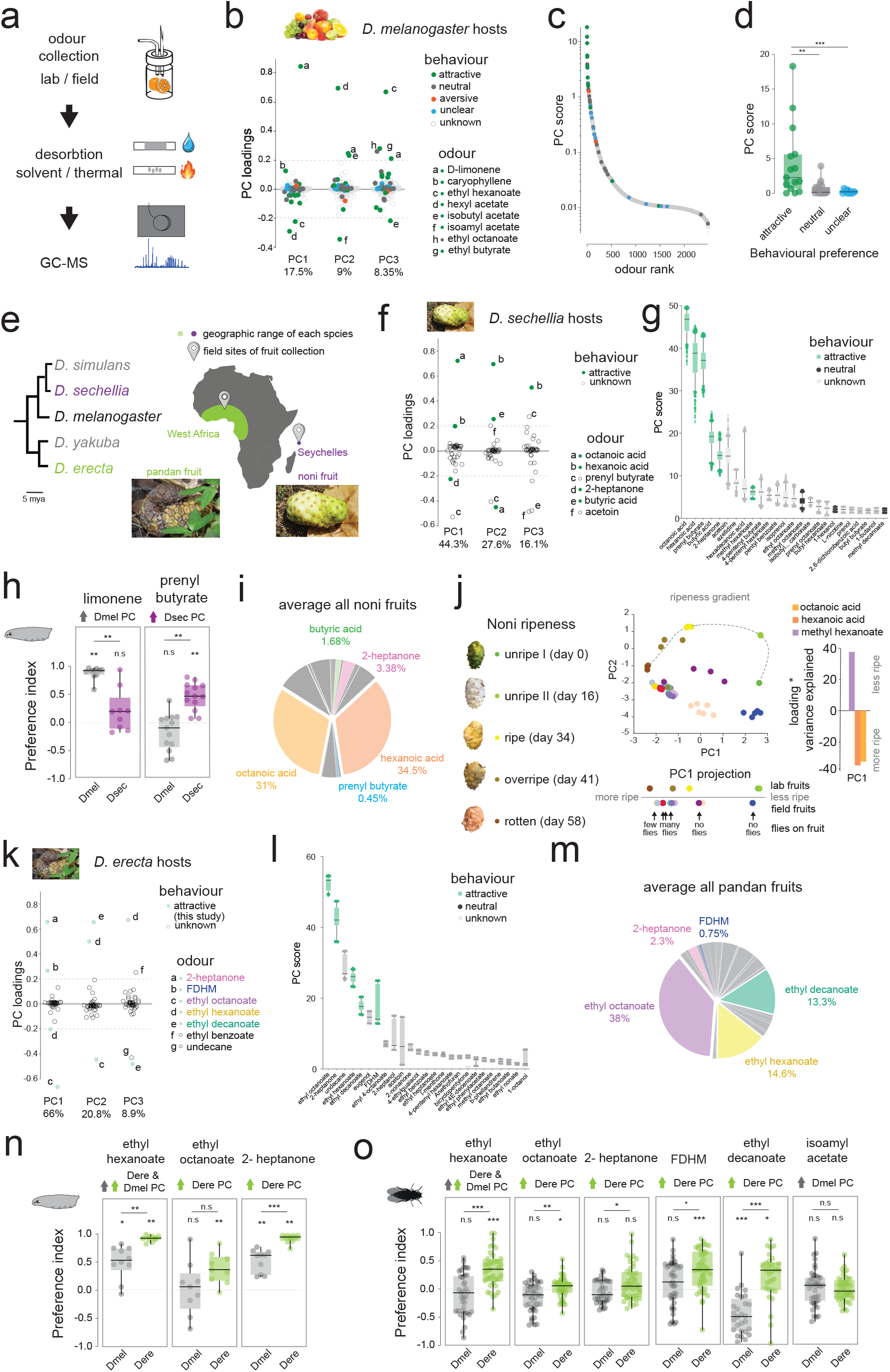
Variance in natural odour blends predicts behaviourally relevant cues across Drosophila species. **a**, Overview of chemical workflow. Host fruit headspace volatiles were collected either from published datasets (of fruits sampled in the laboratory) or from fruits sampled in the field, trapped on sorbent material, desorbed (solvent extraction or thermal desorption), and analysed by GC–MS to obtain relative abundance profiles of volatile compounds (see Methods). **b**, Principal component (PC) loadings for the first three PCs derived from a published dataset^4^ of volatiles emitted by 34 host fruits of *D. melanogaster* (2,491 detected compounds). High-loading compounds dominate variance across fruit odours (values in Table S1). **c**, Behavioural annotation of individual odorants plotted against their PC score (variance-weighted contribution across the first three PCs). Colours indicate behavioural classification from literature: attractive (green), neutral (grey), conflicting (blue), aversive (red) (see Table S2). **d**, Distribution of PC scores grouped by behavioural class. Attractive compounds show higher scores than neutral or conflicting compounds (Mann–Whitney tests; **P = 0.0036, ***P = 0.0007). **e**, Phylogenetic relationships of analysed species and sampling locations. *D. melanogaster*, generalist; *D. sechellia*, noni specialist (Seychelles); *D. erecta*, pandan specialist (West Africa). **f**, PC loadings (first three PCs) from TD–GC–MS analysis of field-collected noni fruits (12 fruits; 38 detected compounds). All values can be found in Table S1. **g**, Robustness analysis of noni PC scores. PCA was repeated 1000 times using random replicate selections per fruit. Points show individual iterations; boxplots show median and interquartile range. Odours coloured by known behavioural effects in *D. sechellia* (attractive, neutral, unknown). **h**, Larval odour preference assays for *D. melanogaster* (grey) and *D. sechellia* (magenta) exposed to limonene and prenyl butyrate. Boxplots show median and interquartile range. Each dot represents a experimental replicate (plate). n = 9 -13. Significance vs neutrality (preference index = 0) and between species were assessed using Wilcoxon tests either one-sample or rank-sum respectively. ** p < 0.005. n.s = non-significant. **i**, Mean relative abundance of identified noni volatiles across field-collected fruits. **j**, Ripeness analysis of noni fruits. PCA combining staged greenhouse fruits^3^ and field fruits shows PC1 corresponds to ripeness stage; fly presence recorded during sampling is indicated below. To the left the loadings of the first three compounds of PC1. **k**, PC loadings (first three PCs) for pandan fruits (8 fruits; 87 detected compounds). **l**, Robustness analysis of pandan PC scores (1000 random replicate iterations as in g). **m**, Mean relative abundance of identified pandan volatiles across field-collected fruits. **n**, Larval odour preference assays comparing *D. melanogaster* and *D. erecta* responses to the indicated volatiles. Boxplots show median and interquartile range. Each dot represents a experimental replica (plate). n = 9 -10. Significance vs neutrality (preference index = 0) and between species were assessed using Wilcoxon tests. ** p < 0.005. n.s = non-significant. **o**, Adult oviposition preference assays for *D. melanogaster* and *D. erecta* to the indicated volatiles. Boxplots show median and interquartile range. Each dot represents a experimental replicate (single fly in a chamber). n = 28-44. Significance vs neutrality (preference index = 0) and between species were assessed using Wilcoxon tests. ** p < 0.005. n.s = non-significant.

We next asked whether this relationship between ecologically informative odour variation and behaviour generalises across species. If olfactory systems evolve to prioritise informative features of their environments, then the odorants that most strongly structure natural variation within a species’ host odour should also be the compounds that drive attraction.

To test this prediction, we examined two non-melanogaster specialist species that have emerged as models for the evolution of odour-guided behaviours^3,13–16^: *D. sechellia*, which feeds almost exclusively on noni fruit^17^, and *D. erecta*, a Pandanus specialist^16,18^ (Fig. 1e). *D. sechellia* provides a well-characterised system with known host cues, allowing validation against established behaviours, whereas the comparatively understudied olfactory ecology of *D. erecta* offers an opportunity to test if our approach can identify novel host-associated cues.

We first tested whether informative variance predicts behaviour in *D. sechellia*. We collected volatiles from noni fruits at different ripening stages and locations in the Seychelles (Fig. 1 e, and Methods), and characterised their chemical profiles using thermal desorption-gas chromatography-mass spectrometry (TD-GC-MS; Fig. 1e, Table S1 and Methods). We analysed these chemical profiles using principal component analysis as above across all noni fruits (Fig. 1f). Furthermore, to test the robustness of our analyses, we collected several replicates per fruit and ran the same analysis iteratively on randomised replicates (Fig. 1g and Methods). Both analyses identified the key components underlying natural variation in noni odours as: octanoic acid, hexanoic acid, 2-heptanone, butyric acid, prenyl butyrate and acetoin (Fig. 1f-g). The first four are already known to play key roles in *D. sechellia* ecology^3,14,19^, again indicating that variance structure recovers behaviourally relevant cues.

This analysis also generated new predictions. Prenyl butyrate ranked amongst the top-scoring odours despite not having been previously implicated in *D. sechellia* sensory ecology. Consistent with our hypothesis, behavioural assays showed that prenyl butyrate is attractive to *D. sechellia* but not to *D. melanogaster* larvae (Fig. 1h), identifying a previously unrecognised host-associated cue. Conversely, limonene - the top-ranked compound in the *D. melanogaster* analysis - attracted *D. melanogaster* but not *D. sechellia* larvae (Fig. 1h). These reciprocal responses underscore the species specificity and predictive power of the approach. Consistent with these results, methyl hexanoate^3^, another ecologically relevant compound for *D. sechellia*, had a high PC score in the noni analysis, whereas three odours reported to be neutral for *D. sechellia*^3^ (1-hexanol, ethyl octanoate and methyl octanoate) showed low scores (Fig. 1g and Table S2).

Together, these findings show that the approach generalises across species and reliably prioritises candidate behaviourally relevant cues. Notably, while some of the identified volatiles are major components of noni fruit, others are only found in relatively low abundance. For example, 2-heptanone is on average 3.38% of all detected noni volatiles, whereas butyric acid accounts for just 1.68%, and prenyl butyrate for only 0.45% (Fig. 1i and Table S3). Thus, behavioural relevance is not simply explained by abundance. Indeed, some of these odorants were originally discovered only through targeted electrophysiology coupled to GC-MS and behavioural assays^14,20^. This underscores that the structure of natural odour variation can be leveraged to prioritise candidate odorants for behavioural and physiological testing.

We next asked which ecological factors underlie natural variation in noni odour blends. We reasoned that fruit ripeness was a strong candidate, as it is known to be an important ecological cue for drosophilids^21–23^. The fruits we collected in the field showed qualitative indicators of ripeness, such as colour and the number of flies present at the time of collection (Fig. 1j),but could not be staged precisely. We therefore used an existing dataset of greenhouse-grown noni fruits that had been accurately staged for ripeness^3^. Analysis of the two integrated datasets revealed that the first principal component corresponded closely with ripeness stage, with greenhouse fruits ordered along PC1 according to ripeness. Importantly, our field collected fruits aligned along the same axis, in positions consistent with their ripeness descriptors (Fig. 1j). The three odorants contributing most strongly to PC1 have all previously been shown to be involved in host seeking and oviposition in *D. sechellia*^*3,14,19*^ (Fig. 1j). Specifically, positive loadings signal less ripeness and are associated with long-range host-seeking^3^, while those odours with negative loadings that signal more ripe fruits have been shown to mediate egg laying^19,24^. This suggests that *D. sechellia* preferentially respond to odours that are most informative about noni fruit ripeness, and exploits the dominant structure of ripeness-related odour variation in noni to guide divergent behaviours: host-seeking vs egg laying.

Finally, we asked whether this framework could predict behaviourally relevant cues in a species with little prior olfactory characterisation. We conducted field work in West Africa, collected volatiles from its host Pandan fruits and characterised their chemical profiles through TD-GC-MS (Fig. 1e, Methods). We identified the top-weighted volatile compounds, dominating variance structure across fruits, as: ethyl octanoate, 2(3H)-furanone, dihydro-3-methyl (FDHM), 2-heptanone, ethyl hexanoate and ethyl decanoate (Fig. 1k-l). Because none of these odorants had previously been linked to *D. erecta* behaviour, they constituted explicit predictions of candidate host-associated cues. Behavioural assays confirmed these predictions. For both adults and larvae, these high-scoring components were more attractive to *D. erecta* than to *D. melanogaster*: most elicited robust attraction and oviposition preference in *D. erecta*, whereas responses in *D. melanogaster* were weak or neutral (Fig. 1n-o). Conversely, isoamyl acetate, identified as informative in the *D. melanogaster* analysis and known to stimulate feeding, led to a subtle, albeit non-significant, oviposition attraction in *D. melanogaster* and was neutral for *D. erecta* (Fig. 1o).

As with our previous analyses, some of the key volatiles for *D. erecta* are found in low abundance, e.g. 2-heptanone and FDHM constitute on average 2.3% and 0.75% of total fruit volatiles (Fig. 1m), reinforcing that behavioural importance is not simply determined by concentration. These results suggest that the *D. erecta* olfactory system, like those of the other species, leverages the information content of individual odour blend components for host-driven behaviours.

Together, our results suggest that host odour-guided behaviours across drosophilids have evolved, at least in part, to exploit the most informative components of natural odour blends. Further, we show that these cues can be predicted directly from environmental statistics. Although PCA has previously been used to separate host identities based on chemical profiles^1,25–27^, we demonstrate a novel use to reveal compounds that guide olfactory decisions in animals, grounded in odour statistics and efficient coding. Drosophilids undoubtedly rely on additional cues not captured by our analyses to navigate the complex chemical environments where they live. Furthermore, different behavioural contexts - such as feeding, oviposition, or larval foraging - may rely on partially distinct odour features^3,4,19,28^. Because our approach is agnostic to behavioural category, it cannot discriminate among these functions. Yet, as suggested by our data, it is likely that these different behaviours are driven, at least in part, by different features of natural odour variation. Therefore, our approach provides a principled starting point for prioritising candidate odorants for targeted testing. More broadly, the ability to predict behaviourally active compounds from a high-dimensional mixture, including low-abundance components, directly from environmental statistics may accelerate discovery in chemical ecology. Such approaches could streamline identification of bioactive volatiles across systems, with potential applications ranging from vector control to agriculture.

## Methods

### Fruit volatile datasets and sampling

For *D. melanogaster* hosts we analysed a published dataset^4^ consisting of headspace odours from 34 fruit species considered potential hosts of *D. melanogaster*, collected in the laboratory using Super-Q® adsorbent filters with hexane solvent elution followed by GC–MS.

For *D. sechellia* hosts, we sampled the fruits of *Morinda citirfolia* trees in the Seychelles, across two of its main islands Mahe and La Digue. For *D. erecta* hosts, we sampled the fruits of *Pandanus candelabrum* in Burkina Faso, West Africa. In both cases, the sampling was done with PDMS tubing as described below.

The ripening analysis of noni fruits was performed on a published dataset of laboratory-staged *M. citrifolia* fruits collected using DVB/carboxen/PDMS SPME fibres followed by GC–MS^3^.

#### Methodological considerations

Passive adsorption onto PDMS tubing does not capture the odour environment identically to insect olfaction. Highly volatile or low-affinity compounds may be under-represented; therefore analyses were interpreted in terms of relative composition across fruits and ripening stages. Additionally, the literature datasets analysed here were collected using different trapping approaches (Super-Q and SPME fibres). These methodological differences limit direct comparison of absolute chemical composition across studies. However, the convergence of predictions across independently collected datasets and multiple sampling chemistries indicates that the variance-based analytical framework is robust to volatile collection methodology.

### Odour extraction of field collected samples

Odour collection and extraction of field collected samples was done according to^29^. Briefly, PDMS tubing (Reichelt Chemietechnik; inner diameter 1.5 mm, outer diameter 2.3 mm) was cut into 5 mm segments and cleaned by immersion in 4:1 acetonitrile:methanol for 3 h, followed by conditioning at 210 °C for 1.5 h (TC-20 tube conditioner, Markes) under a constant nitrogen flow. Tubes were stored in argon-flushed glass vials until use.

For sampling, 3–6 PDMS segments were suspended near each fruit without contacting the surface. Fruit and adsorbents were enclosed in a sealed oven bag and exposed for 30–120 min. Eight pandan fruits and twelve noni fruits were sampled, each PDMS segment was treated as a replicate per fruit. For each fruit, a set of field blanks were collected by manipulating the PDMS segments in the same way but without enclosing a fruit.

After collection, PDMS tubes were frozen and transported to the analytical laboratory for analysis.

### TD–GC–MS analysis

PDMS samples were analysed by thermal desorption gas chromatography–mass spectrometry (TD–GC–MS). Individual PDMS segments were desorbed in a Gerstel thermal desorption unit coupled to a Gerstel CIS-4 cooled injection system and an Agilent 7890A GC with a 5975C inert XL MSD and HP-5MS UI column.

Desorption was performed at 200 °C with cryofocusing at −50 °C. The GC oven program was 40 °C for 3 min, ramp 5 °C min^−1^ to 260 °C (10 min hold), then 5 °C min^−1^ to 280 °C (5 min hold). Mass spectra were acquired in EI mode (70 eV) over m/z 33–500.

### GC–MS curation and compound annotation

Chromatograms were deconvolved using AMDIS (Automated Mass Spectral Deconvolution and Identification System)^30^ and spectra matched to the NIST library. Peaks without confident matches were annotated as unknowns. In parallel, two approaches were used to identify true peaks. MassHunter unknown analysis was used to identify all peaks, while MassHunter Qualitative Analysis was used to inspect blanks and each of the replicates to remove contaminants and column-bleed peaks. XCMS^31^ was used to identify features enriched relative to blanks. Only peaks supported by both approaches were retained. The generated libraries were used to annotate compounds in the final processed chromatograms and extract relative abundances. Compounds not detected were assigned a value of zero. For the field-collected fruit data, to minimise the inclusion of noise-level signals, features with integrated peak areas below 10,000 were excluded prior to downstream analyses, based on inspection of chromatograms and blank samples.

### Principal component analysis

The goal was to identify candidate cues that animals may use for olfactory decisions rather than to reconstruct mixture composition. Absolute GC–MS peak areas do not correspond directly to sensory input, as olfactory systems operate under strong gain control and lateral inhibition, which normalise overall odour intensity and emphasise relative activation patterns across receptors. In addition, recent work has shown that natural odour spaces are structured by the statistics of odorant co-occurrence in natural mixtures rather than by independent variation of individual compounds^2^. We therefore standardized each fruit profile (row-wise z-scoring, where each fruit is a row) to approximate mixture-contrast representations. This transformation normalises abundance across fruits, reduces the influence of dominant compounds and highlights relative deviations within each fruit bouquet. PCA was then performed in MATLAB using pca(), with fruit samples as observations (rows) and odorants as variables (columns).

For comparison, we repeated the analysis using alternative transformations commonly applied to chemical datasets, including standardization of variables across fruits, rather than within each fruit (column-wise z-scoring) and centered log-ratio (CLR) normalization of proportional data^12^. These transformations, which remove within-sample contrast structure, did not equally prioritise behaviourally relevant odorants (see Fig. S1). We note that PCA on proportional data can approximate within-blend prominence when abundance and variance structures align. However, row-zscore normalisation emphasises relative compositional differences, and the results diverge when dominant compounds do not also drive variance across fruits.

For the published *D. melanogaster* host dataset, a single odour profile per fruit type was available and used as input. Field-collected fruits produced multiple chromatographic replicates (3–6 PDMS samples per fruit, see Table S4). For PCA loading plots, replicate chromatograms were averaged so that each fruit contributed one observation. For robustness analyses, each replicate was used separately as detailed below.

#### PC score

Compound contribution to variance structure was quantified using a PC score derived from PCA loadings. Loadings were weighted by the variance explained by each component:

coeffn_{j,i} = coeff_{j,i} × explained_i

The PC score for compound *j* was defined as the sum of absolute weighted loadings across the first three principal components:

PCscore_j = Σ_{i=1..3} |coeffn_{j,i}|

#### Robustness analysis

To evaluate the stability of compound rankings with respect to replicate sampling, PCA was repeated using replicate-level data rather than average fruit profiles. Replicates were treated as nested within fruit and robustness assessed via bootstrap resampling. For each iteration, one replicate chromatogram quantification per fruit was randomly selected. One thousand combinations were generated and duplicate combinations removed. PCA and PC score calculations were repeated for each combination, and PC scores across combinations were visualised as boxplots ordered by median score.

#### Integration of lab and field collected noni fruits

To compare field noni samples with staged laboratory fruits, common odours across both datasets were extracted, and relative abundances and z-scored, calculated. PCA was computed on the laboratory dataset and field samples were z-scored using the laboratory mean and standard deviation and projected into the same PC space.

### Adult behavioural assays

#### Fly stocks and rearing

Oregon-R *D. melanogaster* and wild type *D. erecta* (14021-0224.01) flies were used. Fly stocks were maintained at 25 °C under a 12 h:12 h light:dark cycle on standard cornmeal medium (cornmeal, yeast, glucose, nipagin, and bavistan) supplemented with Drosophila quick mix medium (Blades Biological).

For oviposition conditioning, we followed the protocol by Gou et al.^32^, 0-2 day post-eclosion flies were housed for 5 days in vials containing standard food supplemented with quick mix medium and active yeast paste prepared with 0.5% propionic acid. To promote egg retention, mixed-sex groups were housed at high density. *Drosophila melanogaster* were kept at 35 females and 25 males per vial, whereas *D. erecta* were maintained at 60 females and 25 males to compensate for reduced egg-laying rates and slower larval development. By the end of the conditioning period, the food surface was densely populated with larvae.

#### Oviposition assay

The oviposition assay was adapted from Gou et al.^32^. The base component containing the oviposition substrate was 3D-printed in black PLA with multiple superposed PMMA layers to create the top sections of the chamber and its closing lid, more detail can be found in the GitHub repository (https://github.com/PrietoGodinoLab/High-throughput_drosophila_oviposition_preference_assay). The original design was modified by dividing each arena into two halves, allowing simultaneous testing of 15 individual females per arena.

Flies were cold-anesthetised on ice and a single female was introduced into each chamber. Both species were assayed in parallel under identical environmental conditions.

Odour stimuli were incorporated directly into standard fly food rather than agarose, as preliminary experiments revealed that *D. erecta* deposited insufficient eggs on agarose substrates. Odor concentrations were as shown in Table S5 and chosen based on pilot experiments and previous studies. Approximately 1.5 mL of control food or odour-supplemented food was pipetted into each well, with each well spanning five adjacent egg-laying chambers to create paired-choice conditions. After the assay period, eggs laid on each substrate were counted. Trials in which fewer than five total eggs were laid were excluded from analysis. Oviposition preference was quantified using a preference index calculated as: PI = (a − b)/(a + b) where a and b are eggs on the odour and control sides respectively.

Oviposition assays were conducted at 25 °C and 80% relative humidity in the dark and were performed in a dedicated behavioural room. Assay duration was 24 hours.

### Larval odour preference assays

The larval behavioural assays used the following fly strains: *D. melanogaster* w1118, *D. erecta* white (14021-0224.07), and *D. sechellia* white (14021-0248.30). Eggs were collected on grape-juice agar with yeast. First-instar larvae (0–2 h) were placed at the centre of 10 cm square plates containing 1% agarose. Odours were applied as ten 10 µL droplets onto parafilm strips on opposite sides of the plate. Assays lasted 30 minutes, after this time the preference index was calculated by dividing the arena in six equal sections, three on each side (I, II and III). The number of larva in each section, as well as the number of larva that had reached the odour was counted and the preference index calculated as:

PI = [((4×reached) + (3×zone III odour) + (2×zone II odour) + (zone I odour)) − ((4×control) + (3×zone III control) + (2×zone II control) + (zone I control))]/total number of larvae

Odours were used at 10^-2^ (v/v) dilution in paraffin oil.

### Quantification and statistical analysis

Statistical analyses were performed in R Project. Larval preference comparisons used Mann– Whitney U tests with Benjamini–Hochberg correction where appropriate. Additional tests used for specific figures are described in figure legends. Data and code can be found at: https://github.com/PrietoGodinoLab/fruit_chemical_analysis

**Figure S1.**
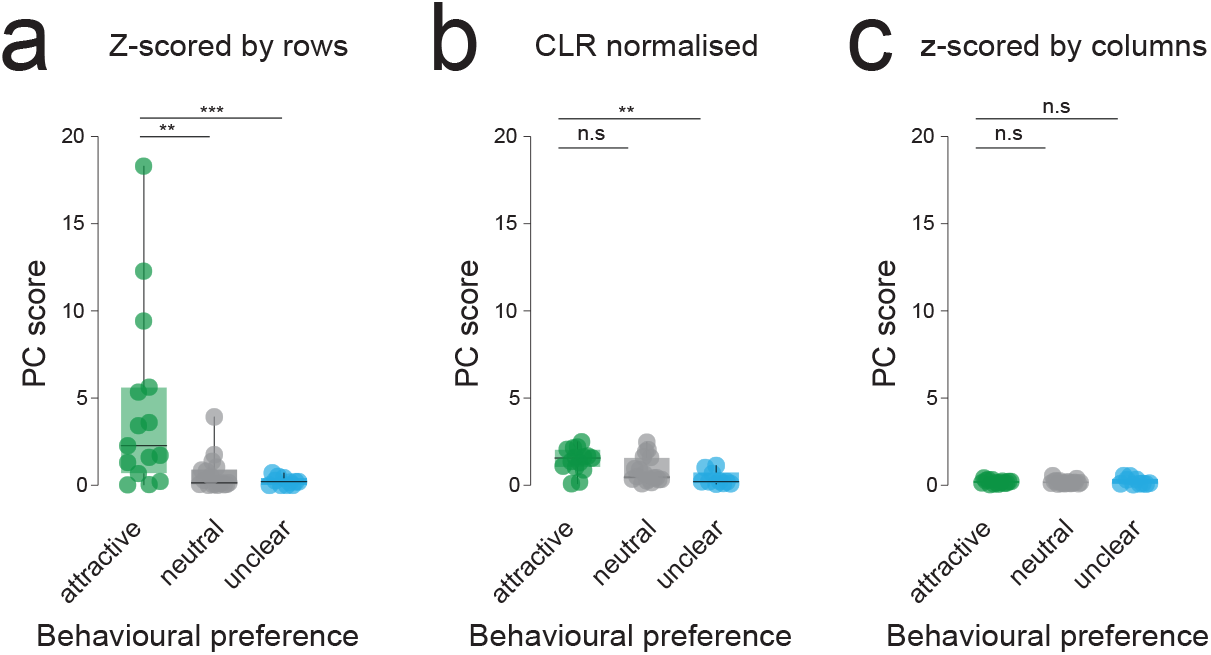
Influence of normalisation on PC score values. Distribution of PC scores grouped by behavioural class based on different normalisations: **a)** z-score across rows (i.e. per fruit), as in Figure 1. **b)** Center-log normalisation, **c)** z-scored across columns (i.e. per odour). Attractive compounds show higher scores than neutral or conflicting compounds (Mann–Whitney tests; **P = 0.0036, ***P = 0.0007).

**Table S1**. Results from the PCA analysis

**Table S2**. Behavioural annotation from the literature

**Table S3**. Compound abundance data

**Table S4**. Chromatogram data for field collected samples.

**Table S5**. Odours used for behavioural assays.

## Acknowledgments

Work in the L.L.P.-G. laboratory is supported by a European Research Council Starting Investigator Grant (802531), an Allen Distinguished Investigator Award by Allen Family Philanthropies, a Human Frontiers Science Grant (RGY0052/2022), a Vallee Scholar Award, a Chan Zuckerberg Collaborative Pairs Project (CP-2-1-Prieto-Godino and by the Francis Crick Institute, which receives its core funding from Cancer Research UK (CC2067), the UK Medical Research Council (CC2067) and the Wellcome Trust (CC2067). We thank Tom Baden for helping with field work in the Seychelles while attracting the majority of the mosquitoes, and comments on the manuscript. We thank Bill Hansson for access to equipment. We thank Albane Imbert from the Crick Making Lab for her role supervising XCF. We thank Jason Somers, Christoph Giez and Andreas Schaefer for comments on the manuscript.

